# PPARD regulation in gastric progenitor cells drives gastric tumorigenesis in mice

**DOI:** 10.1101/218214

**Authors:** Xiangsheng Zuo, Yasunori Deguchi, Weiguo Xu, Daoyan Wei, Rui Tian, Weidong Chen, Micheline J. Moussalli, Yi Liu, Fei Mao, Min Xu, Yaying Yang, Shen Gao, Jonathan C. Jaoude, Fuyao Liu, Mihai Gagea, Russell Broaddus, Keping Xie, Imad Shureiqi

**Author notes:** Correspondence should be addressed to I.S. or X.Z.

## Abstract

Little is known about the cell origin of gastric cancer. Peroxisome proliferator-activated receptor-delta (PPARD) is a druggable ligand-activated nuclear receptor that impacts protumorigenic cellular events. However, PPARD’s role in tumorigenesis, especially gastric tumorigenesis, remains to be defined. We found that targeting PPARD overexpression in murine gastric progenitor cells (GPC), via a villin promoter, spontaneously induced gastric tumorigenesis that progressed to invasive adenocarcinoma. PPARD overexpression in GPC upregulated tumorigenic proinflammatory cytokine and CD44 expression, expanded GPC population *in vivo*, enhanced GPC self-renewal and proliferation in organoid cultures, and endowed these cells with tumorigenic properties. Our findings identify PPARD as a driver of gastric tumorigenesis via GPC transformation.

---

Gastric cancer (GC) is the second leading cause of cancer-related death globally with a 5-year survival rate of less than 25%^1^. Identification of preventive and therapeutic molecular targets in gastric tumorigenesis is needed to improve this poor outcome. Peroxisome proliferator-activated receptor-delta (PPARD) is a ligand-activated nuclear receptor that regulates important molecular cellular events, which strongly impact tumorigenesis (e.g., inflammation, metabolism)^2^. Although PPARD expression is upregulated in many major human cancers^2^, PPARD’s role in tumorigenesis remains controversial^2, 3^. This controversy is primarily based on studies of germ-line PPARD knockout in Apc^min^ mice showing conflicting results regarding the role of PPARD in intestinal tumorigenesis^2^. However, PPARD upregulation is common in human colorectal cancer^2,4^. To study the mechanistic role of PPARD upregulation in colonic tumorigenesis, we generated a mouse model with intestinally targeted PPARD overexpression via the villin-promoter (hereafter designated villin-PPARD mice)^5^.

We serendipitously found that villin-PPARD mice spontaneously developed invasive GC during longitudinal follow-up studies (Figure 1*A-I* and Supplementary Figure 1*A*). Gastric tumorigenesis progressed in age-dependent fashion (Figure 1*I*) from hyperplasia (Figure 1*E*) and low-grade dysplasia (Figure 1*F*) at 25 weeks to high-grade dysplasia (Figure 1*G*) at 35 weeks to large invasive adenocarcinoma at 55 weeks (Figure 1*H*). None of the wild-type (WT) littermates followed concomitantly developed gastric tumors (Figure 1*A-D*, *I* and Supplementary Figure 1*A*). Lesions were initially found in the lesser gastric curvature at 25 weeks (Supplementary Figure 1*A*) and eventually expanded to occupy the whole gastric corpus at 55 weeks (Figure 1*A*). PPARD has been reported to be required for *Helicobacter pylori* infection promotion of gastric epithelial proliferation in rodent and human gastric mucosa^6^. GW501516, a PPARD agonist, promoted DMBA carcinogen-induced squamous GC, a rare form of human GC^7^. These prior reports suggested that PPARD may contribute to GC. These data have remained insufficient to determine the true contribution of PPARD to gastric tumorigenesis, especially the more common adenocarcinoma type. Our new findings demonstrate for the first time that PPARD gene overexpression in villin-positive gastric epithelial cells was sufficient to induce gastric adenocarcinoma. Limited prior studies have shown a vague and questionable relation between PPARD and human GC ^8^. On examining PPARD expression in human GC, we found that PPARD protein expression was higher in epithelial cells of human GC and adjacent tissues than in paired normal gastric tissues (Figure 1*J, K*). Furthermore, PPARD upregulation was associated with lower GC patients’ survival (Supplemental Figure 1*B*). These findings demonstrate the clinical relevance of our mouse data to human GC.

**Figure 1.**
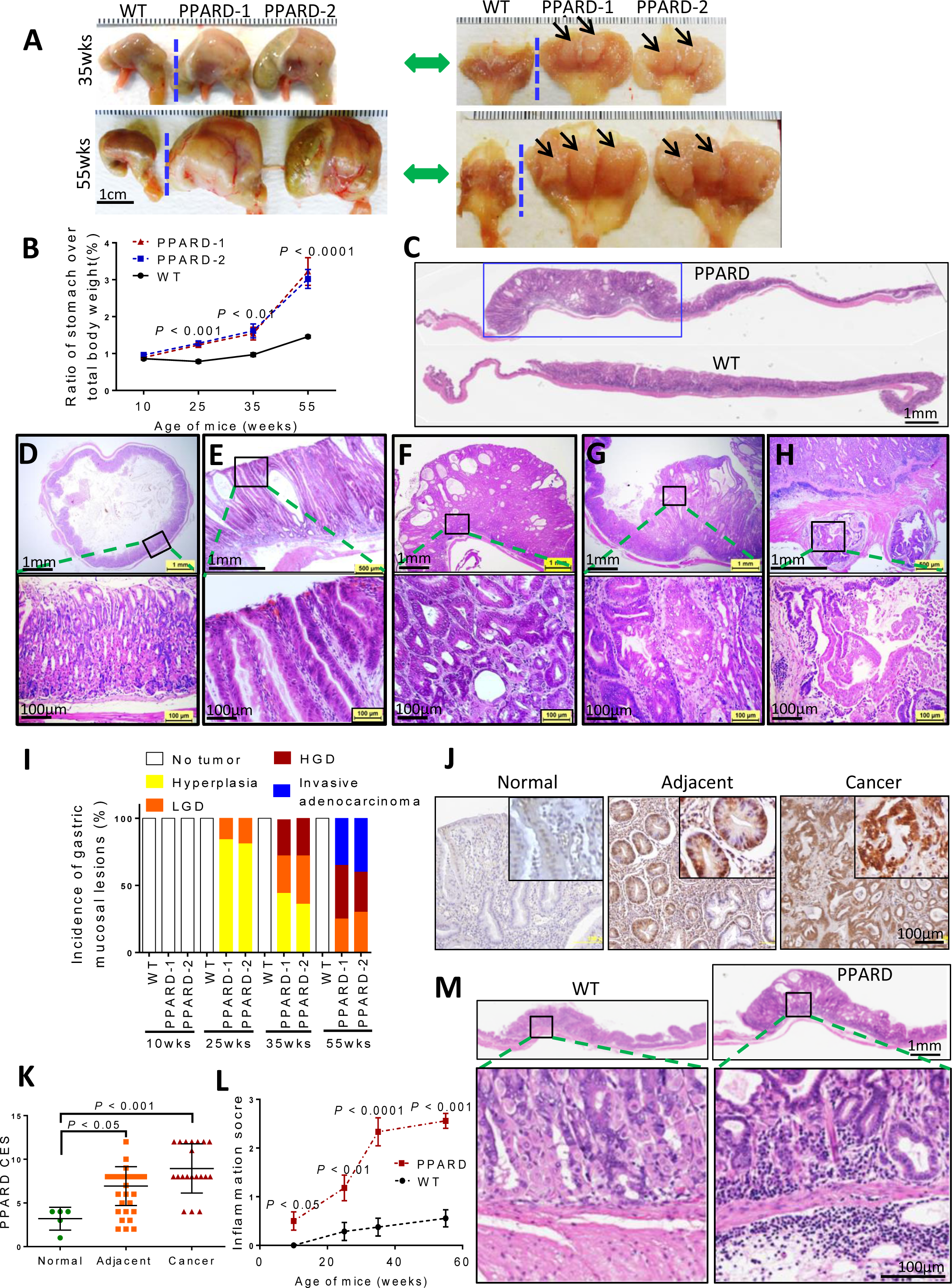
PPARD overexpression in gastric progenitor cells induced large and invasive gastric tumors with inflammation. (*A-I*) Randomly selected villin-PPARD mice from 2 independent founders (PPARD-1 and PPARD-2) and their wild-type (WT) littermates were followed longitudinally and killed at 10, 25, 35, or 55 weeks of age. The stomachs of the mice were removed, weighed, photographed, and examined grossly and histologically. (*A*) Gross examination of the stomachs in PPARD-1 and PPARD-2 mice at the indicated ages. Arrows indicate large nodules within the glandular mucosa in corpuses of stomachs (10- and 25-week findings are shown in Supplementary Figure 1*B*). (*B*) Ratio of mouse stomach weight to mouse body weight in PPARD-1 and PPARD-2 mice and WT littermates at the indicated ages (n =15-20 mice per group). Data are mean ± SEM. ** *P*< 0.01, *** *P*< 0.001, and **** *P*< 0.0001, compared to age-matched WT littermates by 2-way ANOVA. (*C-I*) Histopathological findings of stomachs from PPARD mice. (*C*) Representative photographs of entire gastric mucosa showing the size and extent of lesions (blue rectangle) in PPARD mice in comparison with normal WT littermate mice at age of 35 weeks. (*D-H*) Hematoxylin and eosin (H&E) staining of representative histological features of gastric mucosa from WT mice (normal; *D*) and from PPARD mice (*E-H*) showing a sequence of morphological changes of the stomach from (*E*) marked hyperplasia of glandular epithelium to (*F*) low-grade dysplasia (LGD) and (*G*) high-grade dysplasia (HGD) of glandular epithelium and to (*H*) invasive adenocarcinoma into the muscle layers and lymphatic vessels of stomach wall.(*I*) Incidence of gastric mucosal lesions in PPARD mice according to age. (*J, K*) PPARD protein expression in human gastric cancer. Immunohistochemical staining (IHC) for PPARD was done in stomach adenocarcinoma tissue arrays with self-matched cancer, adjacent tissue, and normal tissue. (*J*) Representative images of IHC staining of the indicated tissues. (*K*) IHC combined expression scores (CES) for human gastric tissue microarrays (n = 26 samples per group). Data are mean ± SEM. * *P*< 0.05, *** *P*< 0.001 by 1-way ANOVA. (*L, M*) Chronic inflammation of gastric lesions of PPARD mice in comparison with stomach of WT littermates at ages from 10 to 55 weeks. (*L*) Comparison of chronic inflammation scores between PPARD and WT littermates at the indicated ages. Data are mean ± SEM. * *P*< 0.05, ** *P*< 0.01, *** *P*< 0.001, and **** *P*< 0.0001 compared to age-matched WT littermates by 2-way ANOVA. (*M*) H&E stained sections of gastric mucosa showing marked transmural infiltration of many lymphocytes in PPARD mice in comparison with normal gastric mucosa of WT littermate mice at the age of 35 weeks.

Chronic inflammation scores of gastric mucosa were significantly higher in villin-PPARD mice than their WT littermates, and these differences increased with age (Figure 1*L, M* and Supplementary Figure 1*C, D*). Progression of lesions in villin-PPARD mice from hyperplasia to invasive adenocarcinoma was associated with increase in chronic inflammation scores (Supplementary Figure 1*D*). These findings agree with the established role of chronic inflammation in gastric tumorigenesis^9^. PPARD overexpression in villin-PPARD mice upregulated the expression of *IL1A*, *TNF-α*, *TLR2*, and *IFNGR1* in gastric mucosa (Supplementary Figure 2*A-D*). These inflammatory cytokine signaling pathways have been implicated in the promotion of gastric tumorigenesis ^10, 11^. Interestingly, treatment of an established human GC cell line (AGS) with IFN-γ (Supplementary Figure 2*E,F*) or a reactive oxygen species inflammatory mediator (H_2_O_2_) (Supplementary Figure 2*G,H*) strongly induced PPARD expression, suggesting a positive feedback loop between PPARD expression and gastric inflammation.

IFN-γ increases proliferation of villin-positive gastric epithelial cells, which have been identified as long-lived quiescent gastric progenitor cells that can generate multilineage cell populations to reconstitute entire gastric glands^12^. Those cells are called villin-positive gastric progenitor cells (VPGPCs). In villin-PPARD mice, PPARD overexpression was initially targeted via the villin promoter to VPGPCs. However, PPARD overexpression subsequently increased villin and CD44 expression in gastric epithelial cells (Figure 2*A-F*) and expanded gastric progenitor cell compartments in gastric crypts of villin-PPARD mice (Figure 2*E*). CD44 is a putative GC stem cell marker that regulates normal and metaplastic gastric epithelial progenitor cell proliferation^13, 14^ and significantly contributes to inflammation promotion of gastric tumorigenesis^11, 15−18^.

**Figure 2.**
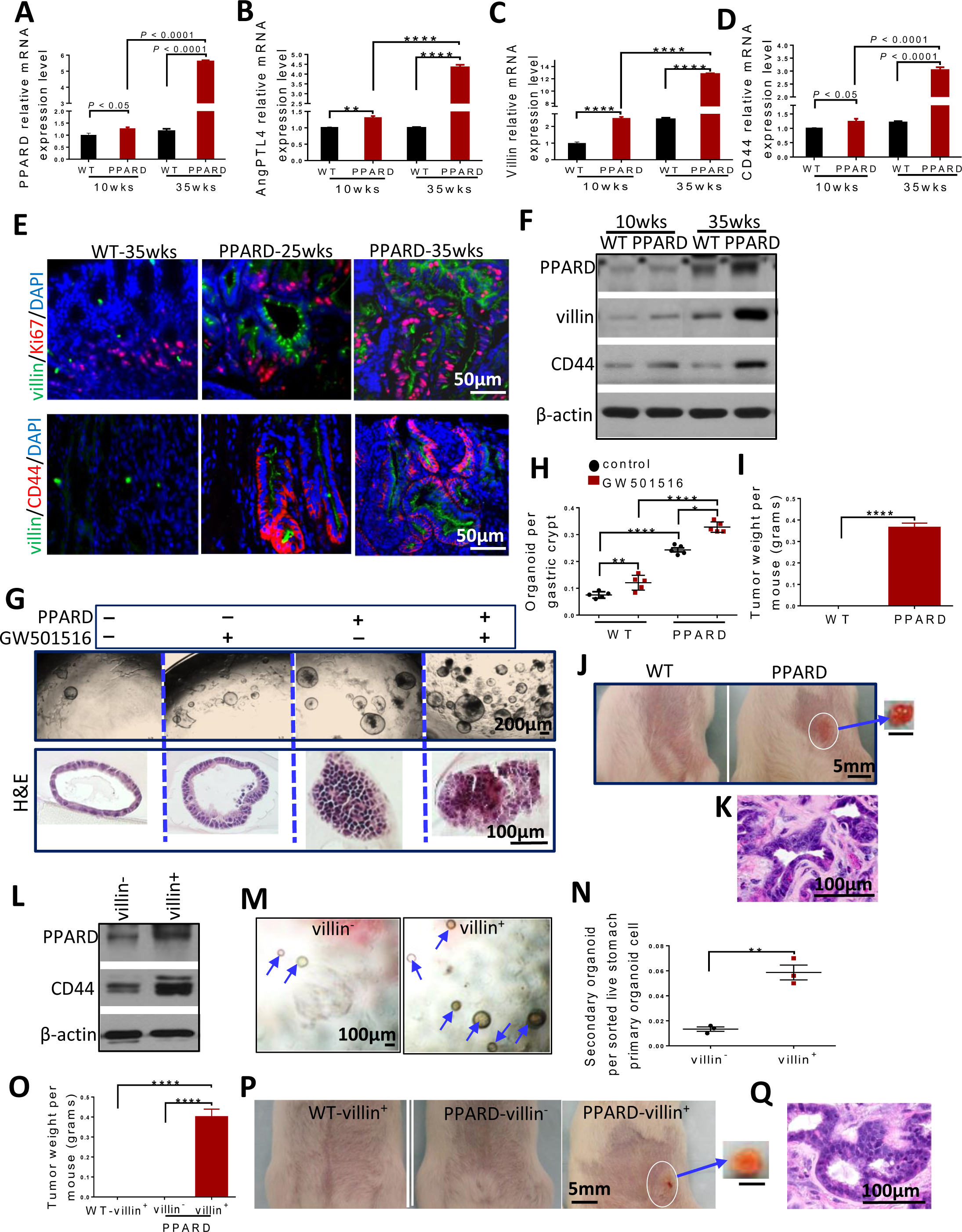
PPARD enhances self-renewal and transforms gastric progenitor cells with tumorigenic properties. (*A-F*) PPARD, villin, and CD44 expression in gastric mucosa in PPARD mice. mRNA expression levels of PPARD (*A*), PPARD target gene ANGPTL4 (*B*), villin (*C*), and CD44 (*D*) of scaped gastric crypts from PPARD mice and wild-type (WT) littermates before (age, 10 weeks) and after (age, 35 weeks) gastric tumor development, measured by quantitative real-time polymerase chain reaction (qRT-PCR). Data are mean ± SD. ** *P*< 0.01, and **** *P*< 0.0001 using 2-way ANOVA. (*E*) Representative images of villin and Ki67, and villin and CD44 expression in gastric mucosa in mice of the indicated genotypes and ages, measured by immunofluorescence staining (n = 3 mice per group). (*F*) Protein expression levels of PPARD, villin, and CD44 of gastric crypts from the same mice described in panels *A-D*, measured by Western blot. (*G, H*) Comparison of organoid-initiating capacity of gastric crypts from 10-week-old PPARD mice and WT littermates fed with GW501516 (10 mg/kg) or control diet (n = 5 mice per group). *(G)*Representative images of primary gastric organoids at day 6 of culture and corresponding representative H&E-stained images. (*H*) Organoid numbers per isolated gastric crypt for the groups described in panel *G*. Data are mean ± SEM.* *P*< 0.05, ** *P*< 0.01, and **** *P*< 0.0001 by 2-way ANOVA. (*I-K*) Tumor formation by primary gastric organoids. Spheroid-derived cells from primary gastric organoids from PPARD and WT littermate mice were subcutaneously injected into syngeneic FVB/N WT mice (n = 5 mice per group). The tumors were assessed 14 days after injection. (*I*) Tumor weights per group. Data are mean ± SEM. **** *P*< 0.0001 by unpaired *t*-test. (*J*) Representative images of the injected mice and isolated tumors. (*K*) Representative H&E-stained image of the tumors described in panels *I, J.*(*L-Q*) Villin-positive and villin-negative cell subpopulations of primary organoid cells from PPARD and WT littermates (n = 3 mice per group) sorted by flow cytometry were harvested for secondary organoid culture. (*L*) PPARD and CD44 expression levels of the sorted villin-positive and villin-negative cells from PPARD mice, measured by Western blot. (*M, N*) Representative images of secondary organoids (*M*) and secondary organoid numbers per sorted living cell (*N*) for the indicated groups from PPARD mice. Data are mean ± SEM. ** *P*< 0.01 by unpaired *t*-test. (*O-Q*) Tumor formation of secondary organoids from the sorted villin-positive and villin-negative cells injected subcutaneously into FVB/N WT mice (n = 3 mice per group). The tumors were assessed 14 days after injection. (*O*) Tumor weights per group. Values are mean ± SEM. **** *P*< 0.0001 using 1-way ANOVA. (*P*) Representative images of the injected mice and isolated tumors. (*Q*) Representative H&E-stained image of the isolated tumor described in panels *O*and *P*.

We therefore examined the effects of PPARD on gastric progenitor cells’ self-renewal capacity and tumorigenicity. PPARD overexpression or activation by PPARD agonist GW501516 treatment markedly increased gastric epithelial crypts-derived spheroid formation in *ex vivo* 3-dimensional gastric organoid cultures (Figure 2*G,H* and Supplementary Figure 3*A-D*), a well-established method to assess cancer stem cell self-renewal. The 3-dimensional cultures of organoids in this experiment were derived from gastric epithelial crypts of villin-PPARD mice at age 10 weeks, before the earliest tumorigenic changes were observed (Supplementary Figure 1*A*). Thus, the observed increases in spheroid formation were driven by PPARD and not secondary to tumorigenesis. PPARD overexpression expanded the CD44-positive and villin-positive cell populations more in villin-PPARD-mouse-derived organoids than in WT-littermate-derived organoids (Supplementary Figure 4*A-C*), further confirming our earlier *in vivo* findings that PPARD promoted progenitor cell self-renewal. Furthermore, PPARD agonist (GW501516) treatment of WT-littermate-derived organoids not only increased spheroid numbers but also produced irregularly shaped spheroid lumens with lining-cell pseudostratification and apoptotic luminal debris, a sign of fast proliferation; in contrast, WT-littermate-derived organoids without PPARD agonist treatment exhibited well-organized circular/oval spheroids lined by a single layer of cuboidal/low columnar epithelium (Figure 2*G,H*). PPARD overexpression in villin-PPARD-mouse-derived organoids produced multilayered, disorganized, irregular epithelial cells that failed to form a lumen, a feature indicative of transformation (Figure 2*G*). Treatment of villin-PPARD-mouse-derived organoids with PPARD agonist GW501516 further increased spheroid numbers (Supplementary Figure 3*A-D*) and produced spheroids with multilayered, irregular cells that appeared to be attempting to form multiple small lumens, resulting in a lumen-in-lumen feature, which is commonly seen in human adenocarcinomas (Figure 2*G*). Furthermore, spheroids derived from villin-PPARD mice, but not their WT littermates, formed tumors when spheroid-derived cells were subcutaneously injected into immunocompetent syngeneic mice (Figure 2*I-K*), thereby demonstrating that PPARD overexpression can enable progenitor cells to acquire a tumorigenic phenotype.

Next, we examined the relatively mechanistic significance of PPARD to the role of gastric progenitor cells in gastric tumorigenesis. Villin-positive cells were selected from organoid-derived cells of villin-PPARD and WT mice by flow cytometry. Compared to villin-negative cells, villin-positive cells from villin-PPARD mice had higher PPARD and CD44 expression (Figure 2*L*) and formed more secondary spheroids in subsequent 3D organoid culture when passaged (Figure 2*M,N*). More importantly, when the sorted organoid cells selected by flow cytometry were subcutaneously injected into immunocompetent syngeneic mice, villin-PPARD-mouse-derived villin-positive organoid cells, but neither WT-mouse-derived villin-positive organoid cells nor villin-PPARD-mouse-derived villin-negative organoid cells, formed tumors in the mice (Figure 2*O-Q*). These latter findings provide strong evidence that PPARD overexpression in villin-positive progenitor cells not only increases progenitor cell self-renewal capacity (Figure 2*L-N*), but also confers on these cells a tumor-initiating capacity (Figure 2*O-Q*).

To further confirm the essential role of PPARD in gastric tumorigenesis, we generated 3 mouse GC cell lines using gastric tumors from 3 villin-PPARD mice at age 55 weeks (Supplementary Figure 5*A*). These villin-PPARD tumor-derived cell lines formed secondary tumors when injected subcutaneously into syngeneic mice (Supplementary Figure 5*B-C*). These cell lines and the secondary tumors expressed the gastric epithelial marker keratin 19 as the gastric epithelial crypts do (Supplementary Figure 5*D*). PPARD downregulation in villin-PPARD-mouse-derived cell lines by lentiviral shRNAs markedly reduced the tumorigenicity of these cell lines when they were subsequently injected subcutaneously into syngeneic mice (Supplementary Figure 5*E-G*), indicating the important contribution of PPARD to the tumorigenicity of these cells. Downregulation of PPARD expression also significantly decreased CD44 expression (Supplementary Figure 5*E*) and attenuated the spheroid-forming ability of these GC cells (Supplementary Figure 5*H,I*), further confirming the mechanistic significance of PPARD to progenitor cell self-renewal.

During the preparation of this manuscript, a report was published showing that PPARD activation enhanced the tumorigenicity of intestinal progenitor cells in APC null mice^19^, which is in agreement with our new findings. However, PPARD’s effects in these cells occurred in association with APC mutation in intestinal cells. APC mutations are a major driver of intestinal tumorigenesis and regulator of stemness; thus, the distinct effects of PPARD are yet to be defined. Our new findings show for the first time in an *in vivo* experimental model that PPARD is sufficient not only to enhance epithelial progenitor cell self-renewal but also, more importantly, to transform these cells so they acquire tumorigenic properties to drive gastric tumorigenesis.

In summary, our results provide the first evidence that PPARD overexpression in progenitor cells is sufficient to induce invasive GC and demonstrate that PPARD can transform gastric progenitor cells. Thus, PPARD is potentially an attractive target for the development of small-molecule PPARD antagonists to prevent and treat GC.

## Supplementary Materials

Methods and any associated references are available in the online version of the paper.

*Note:* Supplementary figures are available in the online version of the paper.

## Footnotes

**Author contributions** I.S. and X.Z. conceived the study, designed the experiments and wrote the paper. X.Z., D.Y., W.X., R.T., W.C., M.J.M., Y.L., F.M., M.X., S.G., J.C.J., and F.L. performed the experiments, and I.S., X.Z. D.Y., and W.X. analyzed the data. W.X., D.Y., Y.Y., M.G., and R.B. performed pathology analyses. D.W. and K.X. provided conceptual feedback.

## REFERENCES

1. Torre LA, et al. CA Cancer J Clin 2015;65:87–108.

2. Xu M, et al. Biochemical Pharmacology 2013;85:607–611.

3. Peters JM, et al. Trends in Endocrinology & Metabolism 2015;26:595–607.

4. Zuo X, et al. JCI Insight 2017;2:e91419.

5. Zuo X, et al. J Natl Cancer Inst 2014;106:dju052.

6. Nagy TA, et al. Gastroenterology 2011;141:553–64.

7. Pollock CB, et al. PPAR Res 2010;2010:571783.

8. Chang YW, et al. Korean J Gastroenterol 2004;43:291–8.

9. Chiba T, et al. Gastroenterology 2012;143:550–63.

10. Castaño-Rodríguez N, et al. Frontiers in Immunology 2014;5.

11. Oshima H, et al. Oncogene 2014;33:3820–9.

12. Qiao XT, et al. Gastroenterology 2007;133:1989–1998.e3.

13. Khurana SS, et al. J Biol Chem 2013;288:16085–97.

14. Takaishi S, et al. 2009;27:1006–20.

15. Bessede E, et al. Oncogene 2015;34:2547–55.

16. Mayer B, et al. The Lancet 1993;342:1019–1022.

17. Ishimoto T, et al. book of Gastroenterology 2015;50:751–757.

18. Khurana SS, et al. book of Biological Chemistry 2013;288:16085–16097.

19. Beyaz S, et al. Nature 2016;531:53–58.

